# Performance Metrics for an Application-driven Selection and Optimization of Psychophysical Sampling Procedures

**DOI:** 10.1101/287904

**Authors:** Mike D. Rinderknecht, Olivier Lambercy, Roger Gassert

## Abstract

When estimating psychometric functions with sampling procedures, psychophysical assessments should be precise and accurate while being as efficient as possible to reduce assessment duration. The estimation performance of sampling procedures is commonly evaluated in computer simulations for single psychometric functions and reported using metrics as a function of number of trials. However, the estimation performance of a sampling procedure may vary for different psychometric functions. Therefore, the results of these type of evaluations may not be generalizable to a heterogeneous population of interest. In addition, the maximum number of trials is often imposed by time restrictions, especially in clinical applications, making trial-based metrics suboptimal. Hence, the benefit of these simulations to select and tune an ideal sampling procedure for a specific application is limited. We suggest to evaluate the estimation performance of sampling procedures in simulations covering the entire range of psychometric functions found in a population of interest, and propose a comprehensive set of performance metrics for a detailed analysis. To illustrate the information gained from these metrics in an application example, six sampling procedures were evaluated in a computer simulation based on prior knowledge on the population distribution and requirements from proprioceptive assessments. The metrics revealed limitations of the sampling procedures, such as inhomogeneous or systematically decreasing performance depending on the psychometric functions, which can inform the tuning process of a sampling procedure. More advanced metrics allowed directly comparing overall performances of different sampling procedures and select the best-suited sampling procedure for the example application. The proposed analysis metrics can be used for any sampling procedure and the estimation of any parameter of a psychometric function, independent of the shape of the psychometric function and of how such a parameter was estimated. This framework should help to accelerate the development process of psychophysical assessments.

## 1 Introduction

Estimating psychometric functions is an important topic, both in basic psychophysics research to investigate mechanisms of sensation and perception, but also in clinical assessments to diagnose sensory deficits, e.g., after neurological injuries. A psychometric function relates physical stimuli to the perception, respectively performance, of the subject in detection and discrimination tasks (Gescheider, 1997). In order to estimate a psychometric function, stimuli of different magnitudes (also referred to as levels) have to be presented to the subject, who then has to rate the stimuli according to the paradigm used in the experiment (e.g., yes-no, same-different, reminder or two-alternative forced choice (2AFC) tasks) (Gescheider, 1985; Macmillan and Douglas Creelman, 2005).

The term *sampling procedure* usually encompasses a set of rules with procedure-specific parameters defining the levels and order of the presented stimuli. There exist many different sampling procedures. The classical sampling procedure is the method of constant stimuli (MOCS) (Gescheider, 1985) presenting stimuli at predefined (i.e., fixed-grid) levels spanning the perception range of interest. While this sampling procedure can be used to obtain the entire shape of the psychometric function, adaptive sampling procedures have been developed to quantify only specific features of a psychometric function (e.g., the perception threshold or slope of a sigmoidal psychometric function) (see Leek (2001); Treutwein (1995) for reviews). These range, among others, from relatively simple staircase (Cornsweet, 1962; Dixon and Mood, 1948; Kaernbach, 1991; Levitt, 1971; Tyrrell and Owens, 1988), heuristic (Findlay, 1978; Taylor and Douglas Creelman, 1967), and stochastic approaches (Kesten, 1958; Robbins and Monro, 1951) to Bayesian (Kontsevich and Tyler, 1999; Prins, 2013; Watson and Pelli, 1983) and maximum-likelihood procedures (Green, 1993; Hall, 1981; Pentland, 1980).

An ideal sampling procedure should be precise and accurate (i.e., present low inherent method variability and be unbiased) to guarantee a high assessment reliability. In addition, sampling procedures should be as efficient as possible (i.e., low number of required trials to achieve a wanted precision) as, especially in clinical settings, time is scarce and costly (e.g., Gresham et al. (1996)). Thus, often, the number of trials is strictly limited due to time constraints or because lengthy experiments could be detrimental for the subject’s attention and lead to mental fatigue (Sullivan and Hedman, 2008). Resulting time-dependent alteration of perception (i.e., drift of psychometric functions) can lead to misestimations of parameters (Doll et al., 2015; Fründ et al., 2011; Leek et al., 1991; Rinderknecht et al., 2018). Moreover, depending on the application scenario, the intra-subject variability may differ and prior knowledge on the distribution of the population of interest may be available or not. As a consequence, different values for the sampling procedure-specific parameters of adaptive sampling procedures defining the stimulus levels to be presented (e.g., around the threshold) may be needed for rapid convergence towards desired features of the psychometric function. Therefore, the question arises how sampling procedures and how sampling procedure-specific parameters should be selected for best performance in a specific application scenario.

As evaluating the performance of sampling procedures through a series of behavioral studies would be too time consuming, computer simulations offer a valuable alternative and powerful tool to simulate psychophysical experiments and to evaluate different sampling procedures and sampling procedure-specific parameters. Besides being used to investigate the process of fitting psychometric functions to psychophysical data (O’Regan and Humbert, 1989; Treutwein and Strasburger, 1999; Prins, 2012; Wichmann and Hill, 2001a,b), computer simulations have been widely used to simulate sampling procedures and quantify their properties, such as the efficiency (Findlay, 1978; Green, 1993; Hall, 1981; Kaernbach, 1991; King-Smith et al., 1994; Madigan and Williams, 1987; Pentland, 1980; Simpson, 1989; Taylor and Douglas Creelman, 1967; Watson and Fitzhugh, 1990). To quantify the efficiency of a sampling procedure, a metric called *sweat factor* was proposed (Taylor and Douglas Creelman, 1967; Taylor, 1971). The sweat factor is defined by the product of the variance of the estimates and the number of trials, in order to evaluate the relative benefit of a longer procedure (i.e., more trials) for a reduced measurement error. Various other approaches and metrics have also been proposed to evaluate the performance of sampling procedures. They commonly include, for example, *mean* or *bias* (e.g., García-Pérez and Alcalá-Quintana (2005); Simpson (1989); Watson and Fitzhugh (1990)), *standard deviation* (e.g., Green (1993); Simpson (1989); Watson and Fitzhugh (1990)), or *settling accuracy* (e.g., Findlay (1978); Pentland (1980)). Others have additionally used *information gain* in bits (Watson and Fitzhugh, 1990) or *percentage usable* (García-Pérez and Alcalá-Quintana, 2005).

However, these performance metrics are commonly used as a function of the number of trials, which may not be the factor which can be acted upon in many application scenarios due to limited time for assessments. When calculated for one given maximum number of trials, the sweat factor’s information content is actually confined to the variability and cannot provide additional information. Furthermore, many simulations sample only one or a very limited number of threshold or slope parameter values (Faes et al., 2007; Findlay, 1978; Green, 1993; Kaernbach, 1991; Madigan and Williams, 1987; Pentland, 1980; Simpson, 1989; Watson and Fitzhugh, 1990). As a matter of fact, the outcome for those metrics may depend on the actual parameter values (e.g., threshold and slope) of the psychometric function to be estimated, and performance may not be homogeneous across this parameter space. Thus, performance results are very likely not representative for other psychometric functions of the population of interest. Instead, constraining the simulations to a specific number of trials given by the requirements of the application and exploring the estimation performance for different psychometric functions covering the entire threshold/slope parameter space of the population would provide relevant insight when selecting and optimizing a sampling procedure for a specific application.

The aim of this paper is to take an application-driven approach and introduce an evaluation framework with a comprehensive set of metrics to analyze psychophysical procedures in terms of threshold and slope estimation performance. We suggest to use error measures widely used in motor control and learning studies (Schmidt and Lee, 2011) in order to describe bias and variability, and introduce *percentage within bounds* (*PCTw*/*iB*) curves, a practical measure depending on desired estimation tolerances which can be directly related to application requirements. Based on this concept, the *normalized area under the curve* (*nAUC*), spanning a surface in a specific threshold-slope parameter space, can be computed. With the *normalized volume under the surface* (*nVUS*) and the inhomogeneity *σ*, we propose measures to compare the performance across different procedures or settings. This framework should facilitate the selection and optimization of sampling procedures for specific applications.

Inspired by a real-world scenario—assessment of pro-prioceptive joint angle difference thresholds using a 2AFC paradigm in a clinical setting—the metrics are illustrated and discussed here on six different procedures using computer simulations: (i) MOCS (Gescheider, 1985), (ii) Weighted Up-Down method (Kaernbach, 1991), (iii) slightly altered Parameter Estimation by Sequential Testing (PEST) (Taylor and Douglas Creelman, 1967; Rinderknecht et al., 2014), (iv–v) standard and accelerated Stochastic Approximation (SA) Staircases (Robbins and Monro, 1951; Kesten, 1958), and (vi) the Bayesian Ψ (PSI) method (Kontsevich and Tyler, 1999).

## 2 Definition and parameters of the psychometric function

In the present work, the perception models consisted of psychometric functions *ψ*(*x*) defining the proportion of correct responses at different stimulus levels *x*:

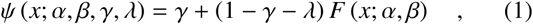

with the generic threshold parameter *α*, generic slope parameter *β*, guessing rate *γ*, and lapse rate *λ* (taking into account stimulus-independent errors, or “lapses”). The guess rate *γ* depends on the psychophysical paradigm (e.g., yes-no: *γ* = 0, 2AFC: *γ* = 0.5). The lapse rate *λ* is often set to 0, to limit the complexity of computer simulations. For the generic sigmoid function *F*(*x*; *α*, *β*), a cumulative normal function *F*_*Gauss*_(*x*; *µ*, *σ*) with a mean µ and standard deviation *σ* according to the following equation was chosen:

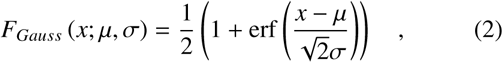

where erf(*x*) is the standard definition of the error function:

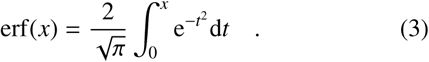

As a consequence of this choice and a lapse rate *λ* of 0, the generic threshold parameter *α* corresponds directly to the inflection point *µ* and the threshold, which is defined at *x*_*T*_ = *ψ*^−1^(*P*_*t*_), being the target probability or proportion of correct responses. The generic slope parameter *β* is inversely proportional to the “spread” (i.e., standard deviation *σ* at this point). In order to have comparable values across different studies using various analytic functions, it has been recommended to use the maximum actual slope *Slope*_*α*_, instead of the slope parameter *β* depending on the type of cumulative distribution. This is achieved by taking the first derivative *d*ψ**/*dx*|_*x*=*α*_ (Strasburger, 2001) according to:

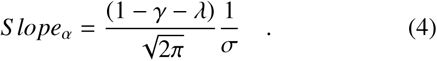

The parameters and psychometric functions are illustrated in Figure 1.

**Figure 1.**
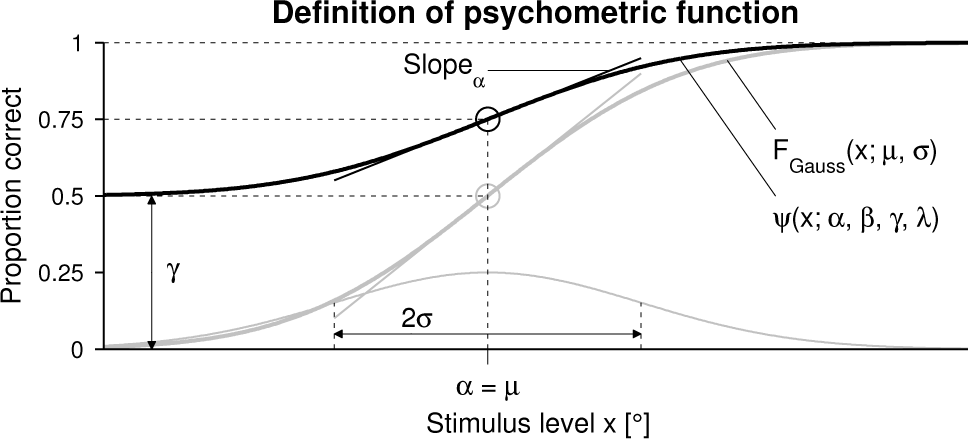
Psychometric function. Illustration of the psychometric function *ψ*(*x*;*α*, *β*, *γ*, *λ*) (bold black sigmoid) and related parameters. The underlying cumulative normal function *F_Gauss_*(*x*;*µ*, *σ*) (bold gray sigmoid) and its underlying normal probability density function (gray) are also indicated. The slopes are indicated at the inflection points in the same color of the corresponding sigmoids. Note that the psychometric function is illustrated for the present application of proprioceptive joint angle assessments using a 2AFC paradigm, where the stimulus level *x* corresponds to the angular difference between two stimuli to be distinguished in a trial in degrees (°). In this application, *γ* was set to 0.5, *λ* to 0, and the difference threshold was defined at *x*_*T*_ = *ψ*^−1^(*P*_*t*_), with *P_t_* = 0.75

## 3 Performance metrics for psychophysical procedures

In order to quantify the estimation performance of a sampling procedure, the “real” value of the parameter in question (i.e., in this case the threshold) of the psychometric function to be estimated should be known. This is where computers simulations come in useful, as the “real” psychometric function can be modeled and is known. Furthermore, the psychophysical experiment using the sampling procedure should ideally be simulated multiple times to obtain a distribution of estimates for high statistical power. To better distinguish between the “real” psychometric function to be estimated and the psychometric function fitted to the data provided by a (simulated) experiment, the first is referred to as template 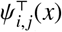 (the symbol Τ denotes a template, and indices *i*, *j* the combination of threshold *α* and slope *Slope*_*α*_ values).

The most elementary analysis consists of quantifying estimation bias (accuracy) and estimation variability (precision). The following nomenclature of constant errors (*CE* = average signed errors) and variable errors (*VE* = standard deviations of errors) was used. This nomenclature is commonly used in motor control and learning (Schutz and Roy, 1973). In order to evaluate the estimation performance of sampling procedure, the threshold estimation error is computed by subtracting the threshold of the template 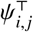 from the threshold of the estimated psychometric function. A positive error represents an overestimation (i.e., larger threshold or larger slope estimates compared to the template), whereas a negative error represents an underestimation (i.e. smaller threshold or smaller slope estimates). Before calculating the slope estimation error in the same way, an arctan(*Slope*_*α*_)/(*π*/2) transform was applied (Figure 2). This transformed the slope space from [0, inf) to [0, 1] in arbitrary units (a.u.), where zero and one corresponded to a completely flat psychometric function and a perfect step function, respectively. Without applying this transform first, slope errors around large slopes (i.e., steep psychometric functions) diverge towards infinity, despite the psychometric functions looking almost identically. Calculating and plotting the *CE* and *VE* for a fine, two dimensional grid of threshold and slope parameter values allows to identify potential zones in the threshold-slope space, which may suffer from poorer estimation performance. This may provide insight on how sampling procedure-specific parameters could be tuned for the psychophysical assessment application.

**Figure 2.**
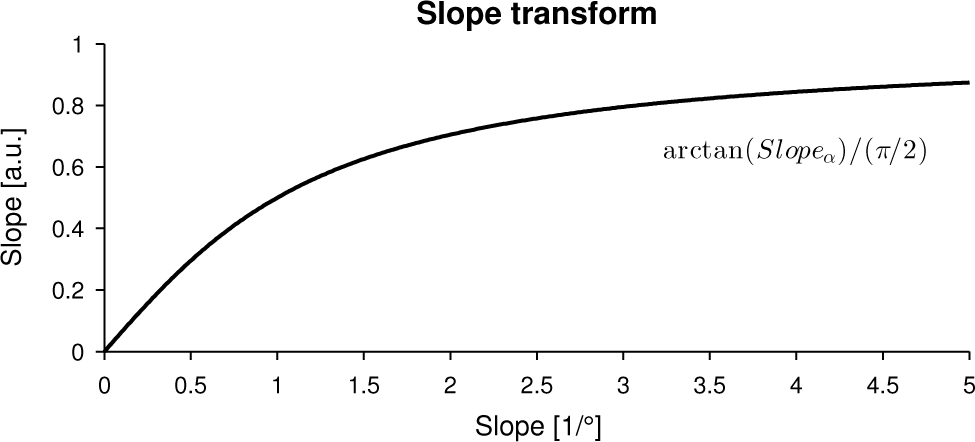
Slope transform. Function transforming the actual slope in stimulus units (in the present application of proprioceptive joint angle assessments in (1/°)) to arbitrary units (a.u.) within the range [0, 1]. This transformation prior to the calculation of the performance metrics is essential to make the estimation errors comparable for different slope values.

Since the absolute errors (*AE* = average absolute errors) are a complex combination of *CE* and *VE* and can be predicted from them (Schutz and Roy, 1973), direct examination of the *AE* becomes superfluous. However, it can be used in an application-driven approach to develop other higher level metrics building upon it. As the required precision depends on the application, it can be useful to describe the probability of a resulting estimation lying within an interval. Thus, the performance of a procedure could be expressed as the percentage of simulation results of threshold and slope estimates lying within a tolerance interval around the template values, respectively, as a function of the interval size. This *percentage within bounds* (*PCTw*/*iB*) function is related to absolute errors as illustrated in Figure 3 (A)–(C). A faster-rising *PCTw*/*iB* function would correspond to a stimulus selection method with higher performance. This metric would allows selecting the optimal sampling procedure given a required maximal error. Note that this metric takes into account both accuracy and precision, but provides more information relevant to the application compared to the elementary absolute error.

**Figure 3.**
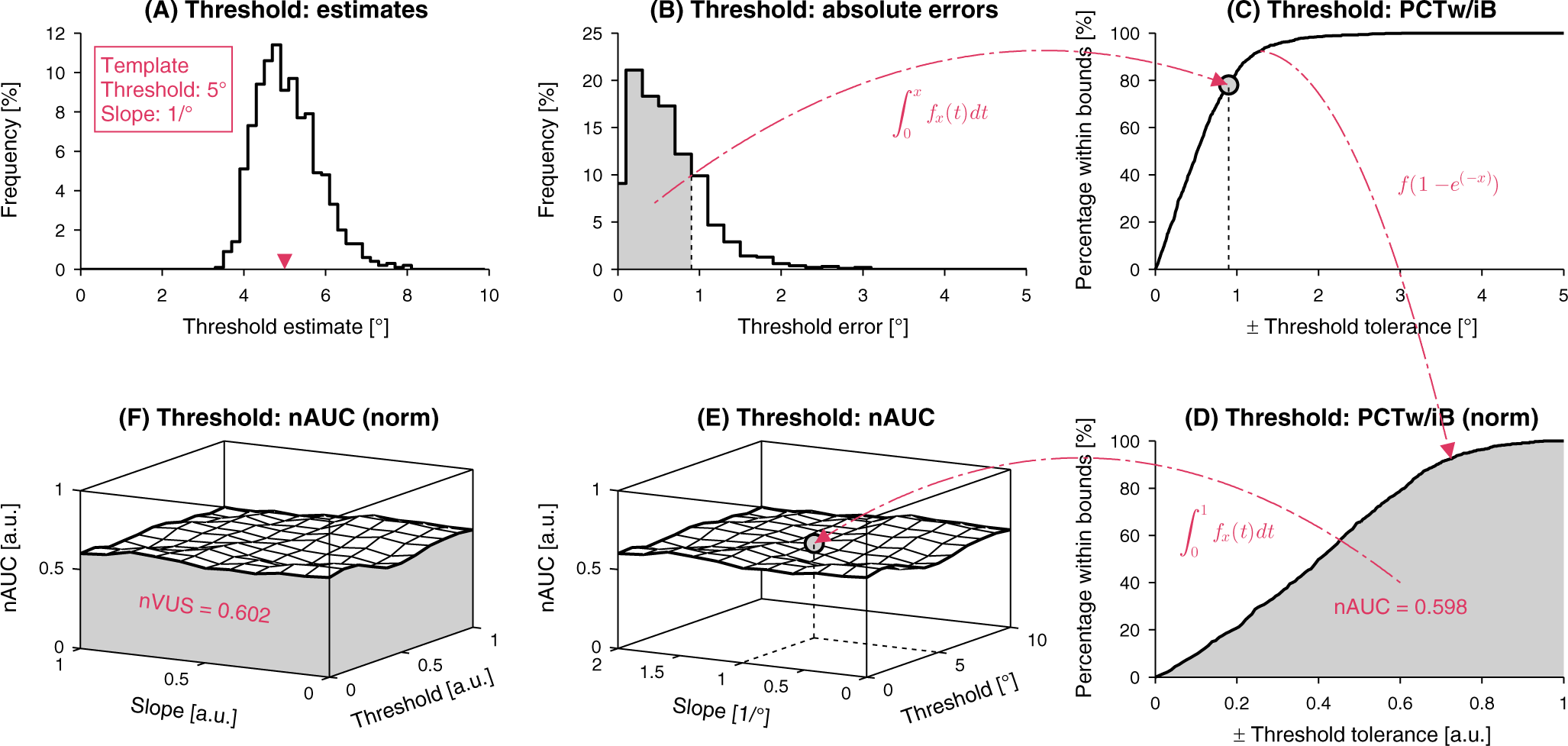
Explanation of performance metrics to characterize psychophysical sampling procedures. **(A)** Distribution of threshold estimates for a given template. **(B)** Distribution of the absolute errors. **(C)** *Percentage within bounds* (*PCTw*/*iB*) as a function of the tolerance interval. Each point of this function is generated by integrating the area of the absolute error distribution from zero to the respective error, respectively tolerance, value. **(D)** After a nonlinear transform of the *x* axis to [0, 1], the *normalized area under the curve* (*nAUC*) can be calculated (1 corresponding to perfect estimation for this template). **(E)** When the *nAUC* is computed for different templates within a threshold and slope parameter space, the performance can be visualized as a surface in a three-dimensional space. **(F)** After normalizing the parameter space axes to [0, 1], the *normalized volume under the surface* (*nVUS*) can be calculated (1 corresponding to perfect estimation across all templates). Note that the metrics are illustrated for the present application of proprioceptive joint angle assessments using a 2AFC paradigm, for a psychometric function template 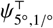.

Similar to the receiver operating characteristic curve, the performance of the sampling procedure can be quantified by the area under the curve (AUC) (i.e., definite integral under the *PCTw*/*iB*-curve). Since the AUC is not bounded in the case of the threshold and could not be calculated without defining an arbitrary upper bound on the tolerance interval size, a nonlinear transform 1 − *e*^(−*x*)^ is applied to the tolerance interval axis *x* of the *PCTw*/*iB*-curve. With this transform, the positive semi-infinite support [0, inf) is transformed to [0, 1] and the *normalized AUC* (*nAUC*) ∈ [0, 1] can be calculated (Figure 3 (D)). Due to the transform applied to the slopes before calculating the errors, the tolerance interval axis is already bounded and normalized to [0, 1]. Therefore, this nonlinear transform is not required anymore and the *nAUC* can be calculated directly for the slopes. The precision of *nAUC* can be improved by increasing the number of simulated runs for each template 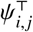. A repeatedly perfect estimation of the threshold or slope parameter would result in a *nAUC* of one. This metric takes all estimates into account without having to calculate the parametric statistics, such as the arithmetic mean for the average absolute error. This metric is still template-dependent and can be used for more high-level metrics quantifying performance over the complete threshold and slope parameter space.

The performance of a procedure may vary depending on the threshold and slope of the psychometric function in question. Therefore, the *PCTw*/*iB* and corresponding *nAUC* for the threshold and slope estimation have to be calculated for each template 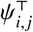. The *nAUC* can be visualized as a surface in a three-dimensional space (Figure 3 (E)). For a given threshold and slope parameter space the *normalized volume under the surface* (*nVUS*) can be calculated after linearly normalizing the threshold and slope axes to [0, 1] (Figure 3 (F)). As a result, the *nVUS* is also ϵ [0, 1] and can be used to compare different procedures or method settings, as long as the same application-dependent parameter space range is used. The accuracy of *nVUS* can be improved by simulating a denser grid of templates 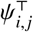. A *nVUS* of one corresponds to perfect estimations for all threshold and slope combinations within the evaluated parameter space. This metric can be used for an overall performance comparison across sampling procedures.

In addition to the *nVUS*, the performance variability can be evaluated: The inhomogeneity of the *nAUC* across the parameter space can be described by calculating the standard deviation *σ* of all the *nAUC* values for the different templates 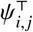. This parameter *σ* should not be confused with the parameter of *F*(*x; µ*, *σ*). To calculate the standard deviation, each axis of the parameter space should be linearly sampled to avoid bias towards *nAUC* values where the sampling density is higher. In case the sampling of the simulated parameter space is not linear along one or both axes, the *nAUC* surface has to be resampled and interpolated along the axes in question.

To have an overall performance measure for each threshold and slope estimate as a function of the accepted estimation tolerance, for each tolerance interval the *PCTw*/*iB*_±*Tol*_ can be presented as a surface for the parameter space, similar to the *nAUC*. From this surface, the overall tolerance-dependent performance can be calculated, as done for the *nVUS* and *σ*.

## 4 Computer simulations

In order to illustrate and discuss the performance evaluation metrics described in the previous section for different procedures, a computer simulation based on a concrete application example was implemented.

### 4.1 Application example: Assessment of proprioceptive difference thresholds

Accurate and sensitive assessments of proprioception may be used for diagnosis, prognosis and treatment planning (Pumpa et al., 2015) for patients with somatosensory deficits affecting the upper limbs (e.g., after neurological injuries and diseases). However, clinical assessments, such as the up-down finger proprioception test (Lincoln et al., 1991), provide mostly dichotomous or ordinal scales and may thus be used for screening, but not for assessing functional improvements (Hillier et al., 2015). The combination of psychophysical procedures to estimate psychometric functions with robotic technology would offer more reproducible assessments with a higher resolution. There have been few studies exploring this approach for the assessment of the upper limb (Brewer et al., 2005; Tan et al., 2007; Lambercy et al., 2011; Rinderknecht et al., 2014; Simo et al., 2014; Elangovan et al., 2014; Cappello et al., 2015). So far MOCS has been predominantly used, with experiments typically lasting about 45 min (Brewer et al., 2005; Tan et al., 2007; Lambercy et al., 2011; Simo et al., 2014). However, in order to achieve clinical utility, the number of trials and the assessment duration should be reduced to below 15 min without compromising the quality of the outcome measures. This may be achieved by using adaptive procedures (e.g., as shown in a pilot study comparing MOCS and the adaptive PEST experimentally (Rinderknecht et al., 2014)).

The problem with such experimental validations is that the “real” psychometric functions are unknown. Therefore, the estimation performance and efficiency of the sampling procedures cannot be directly quantified. This can be addressed in computer simulations, where the actual psychometric function templates are known. Moreover, this allows for optimization of sampling procedure-specific parameter settings for a specific threshold and slope range of the population of interest. So far, experimental results on the difference threshold of angular joint position revealed values ranging from 1 to 5°, approximately, for elbow, wrist, and finger joints in healthy subjects (Tan et al., 1994; Brewer et al., 2005; Tan et al., 2007; Lambercy et al., 2011; Rinderknecht et al., 2014; Elangovan et al., 2014; Cappello et al., 2015). However, in patients with proprioceptive deficits those values are higher and may go up to around 10° (Rinderknecht et al., submitted).

The quality of estimation of difference thresholds can be affected by the used experimental paradigm. While psychophysical experiments based on paradigms such as yes-no, reminder, and same-different can provide quantitative results, they are contaminated by effects of the decision criterion (i.e., response bias) (Gescheider, 1985; Macmillan and Douglas Creelman, 2005). Despite some literature claiming that the two-alternative forced-choice (2AFC) paradigm requires a two-to three-fold number of trials for a given precision compared to the Yes-No paradigm (McKee et al., 1985; King-Smith et al., 1994), 2AFC addresses the previously mentioned limitations, as it is expected to be a more sensitive, more objective and almost bias-free alternative (Macmillan and Douglas Creelman, 2005).

Assessing proprioception at a specific joint using the 2AFC paradigm requires a two-interval design: two different stimuli (i.e., two flexion or extension movements are consecutively presented) before the subject rates which angle, respectively movement from a reference position, was larger. In the present work the difference between the two angles of one trial is referred to as level *x*. Prior work showed that one trial including response time lasts about 15 s (Rinderknecht et al., 2014). Thus within an acceptable assessment duration of 15 min around 60 trials can be administered.

These criteria set the stage for the following computer simulations, illustrating the proposed evaluation metrics for a real-world application example.

### 4.2 Methods

#### 4.2.1 Templates

To cover the full population, a set of 240 subject templates 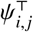 was created with 40 different thresholds linearly spaced within the range [0.25°, 10°] (index *i*) and slopes *Slope*_*α*_ 0.0625, 0.125, 0.25, 0.5, 1, 2 /° (index *j*). The guess rate *γ* was 0.5 and the lapse rate *λ* was zero for all templates. As an example, the template 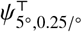 would have a threshold of 5° and a slope *Slope*_*α*_ of 0.25/°. Examples of modeled psychometric functions with a threshold of 5° and different slopes are visualized in Figure 4.

**Figure 4.**
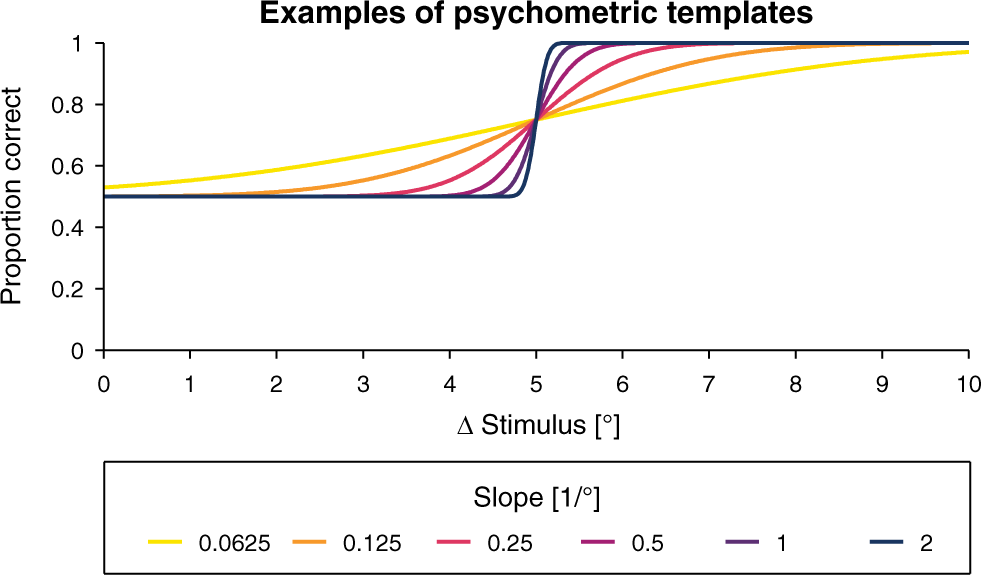
Examples of psychometric templates. Examples of psychometric templates 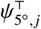 with a threshold of 5° and different slopes used in the computer simulations.

#### 4.2.2 Simulation process

The response of a simulated subject for a given stimulus level *x* was generated by comparing a randomly generated number between 0 and 1 to 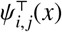. A larger random number compared to 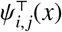 corresponded to a false response, and a smaller random number compared to 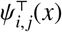 to a correct response.

Six different sampling procedures were simulated. A brief explanation of the sampling procedures and follows in the section below, and the used sampling procedure-specific parameter values are reported. The parameter values were empirically chosen based on experience and reasonable outcomes for this application and population. Thus, we do not claim that the parameter values result in optimal performance for each sampling procedure. Each sampling procedures was simulated for a length of 60 trials, and each template was simulated 1000 times for each psychophysical procedure, leading to a total of 6 × 240000 simulated psychophysical experiments.

Even if some adaptive methods directly provide a threshold estimate, they may not always reach convergence within the given maximum number of trials and therefore threshold estimates may not be very accurate. Thus, the threshold and the slope parameters were estimated by fitting *ψ*(*x*) to the proportion of correct responses at stimulus levels *x* using a Maximum Likelihood criterion implemented in the Palamedes MATLAB routines (Prins and Kingdom, 2009). Moreover, this way the parameter estimation process was identical for all examined methods. The computer simulations, as well as the following data analysis, were performed in MATLAB R2014a (MathWorks, Natick, MA, USA).

#### 4.2.3 Sampling procedures

##### MOCS

The method of constant stimuli (Gescheider, 1985) is the simplest sampling procedure, where a set of stimulus levels is predefined and presented multiple times in random order. The set of stimulus levels used in this simulation consisted of 12 levels spaced equally ∈ [0.75°, 9°]. Each level was presented 5 times.

##### Weighted Up-Down

In contrast to the Simple Up-Down (Dixon and Mood, 1948) and the Transformed Up-Down (Levitt, 1971) methods, the Weighted Up-Down method proposed by Kaernbach (Kaernbach, 1991) can converge to any desired point on the psychometric function using the equilibrium condition *step_up_ P*_*t*_ = *step*_*down*_ (1 − *P*_*t*_) for the convergence point *x*_*t*_ = *ψ*^−1^(*P*_*t*_). Thus, for a target probability *P*_*t*_ = 75%, the ratio *step*_*up*_/*step*_*down*_ results in ^1^/_3_. Each correct or incorrect response leads to a decrease, respectively increase, of the stimulus level according to the following rules:

- −3 *step*_*unit*_ after 3 correct responses,
- +1 *step*_*unit*_ after 2 correct and 1 incorrect response,
- +2 *step*_*unit*_ after 1 correct and 1 incorrect response,
- +3 *step*_*unit*_ after 1 incorrect response,

where *step*_*unit*_ was 0.5° and the start level *x*_0_ was 5.5°.

##### PEST (log)

The adaptive procedure called Parameter Estimation by Sequential Testing was introduced by Taylor and Douglas Creelman (1967). The desired convergence point (percentage of correct responses) can be selected, no prior assumptions on the subject’s psychometric function are required, and it is based on a set of heuristic rules defining step sizes as follows:

- The step size is halved on every direction reversal.
- The first and second step in the same direction are of same size.
- The third step is double the second if the step immediately preceding the last reversal resulted from a doubling, or same otherwise.
- The fourth and additional steps in the same direction are the double of their predecessor.

According to the Wald sequential likelihood-ratio test, the level is maintained if *N*_*correct*_ ∈ (*N*_*total*_ × *P*_*t*_ ± *W*) and changed otherwise. *N*_*correct*_ and *N*_*total*_ correspond to the number of correctly responded trials and the total number of subsequent trials at the same level. A value of *P*_*t*_ = 0.75 leads to a convergence towards 75% correct responses in a 2AFC experiment. The deviation limit *W* of the sequential test was set to *W* = 1, as suggested in (Taylor and Douglas Creelman, 1967). This parameter defines the trade-off between quick (highly dynamic behavior of PEST) and powerful (slower level changes but with higher statistical confidence) decisions. PEST requires two starting parameters: the start level *x*_0_ and the *step*_*start*_, which were set to 5.5° and 2°, respectively. Despite PEST having three termination conditions in addition to a total maximum number of trials (i.e., maximum number of consecutive trials at the same level *x* or a step below a minimum threshold *step*_*min*_), they were not used in the present simulations to reach always 60 trials.

PEST is often used in psychoacoustic experiments where auditory stimulus levels are given in dB (Taylor and Douglas Creelman, 1967; Hall, 1981; Leek, 2001). However, when PEST is applied to an experiment estimating a joint angle difference threshold, zero crossings of the level *x* when continuously decreasing the level leads to potential problems and undesired behavior of the algorithm (convergence towards the upper or lower difference threshold, and reduction of efficiency through temporally divergence from the threshold). To address these issues, a logarithmic mapping *f*: Stimulus → PEST, *f* (*x*) = log *x* between the stimulus domain in degrees and the PEST domain was introduced (Rinderknecht et al., 2014). Consequently, the stimulus levels always remain positive, even if the PEST level performs zero crossings. Because the mapping depends on absolute values, the mapping functions *f* (*x*) and *f* ^−1^(*x*) cannot be directly applied to a step. Instead, a step has to be regarded as a vector and the mapping functions have to be applied to the initial and terminal points separately, after which the two values are subtracted to define the step length in the specific domain (i.e., Stimulus- or PEST-domain).

##### SA Staircases (standard)

The standard Stochastic Approximation (SA) Staircases (Robbins and Monro, 1951) can converge to any target probability *P*_*t*_ using asymmetric upward and downward steps. The step size is defined by the following rule and decreases with the number of trials *n*:

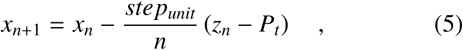

where *z*_*n*_ is the binary response (0 incorrect, 1 correct) at trial *n*. The start level *x*_0_ and *step*_*unit*_ were set to 5.5° and 4°, respectively, with *P*_*t*_ = 0.75.

##### SA Staircases (accelerated)

Kesten proposed an accelerated version of the Stochastic Approximation (SA) Staircases (Kesten, 1958). The first two trials of the procedure are identical to the standard SA Staircase. For the subsequent trials (*n* > 2) the step size is changed only the when response changes according to the following rule:

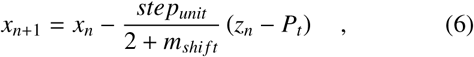

where *m*_*shift*_ is the number of shifts in response category (i.e., reversals). The start level *x*_0_ and *step*_*unit*_ were set to 5.5° and 4°, respectively, with *P*_*t*_ = 0.75.

##### PSI method

The Ψ method was developed by Kontsevich and Tyler (1999) to estimate the threshold and slope: The posterior probability distribution across the two-dimensional space (threshold and slope) of psychometric functions is updated following Bayes’ rule. Subsequently, the psychometric function is estimated by computing the mean of the posterior probability distribution. The next level is defined by a one-step ahead minimum search of an entropy-based cost function in order to optimize information gain. The detailed equations are presented in (Kontsevich and Tyler, 1999). The threshold grid used in the simulations was [0.25°, 10°] in steps of 0.125°. The slope (*Slope*_*α*_) grid consisted of 12 logarithmically spaced values from 0.0625/° to 2/°. The guessing rate γ and lapse rate *λ* were set to 0.5 and 0. Levels were restricted to *x* ∈ [0.25°, 10°] in steps of 0.125°.

### 4.3 Results

#### 4.3.1 Example sequences

A set of procedure sequence examples (from one simulation run) for the six different tested procedures illustrating stimulus placement and adaptiveness (where applicable) are shown in Figure 5 in combination with the template function 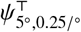 and the resulting estimated psychometric function.

**Figure 5.**
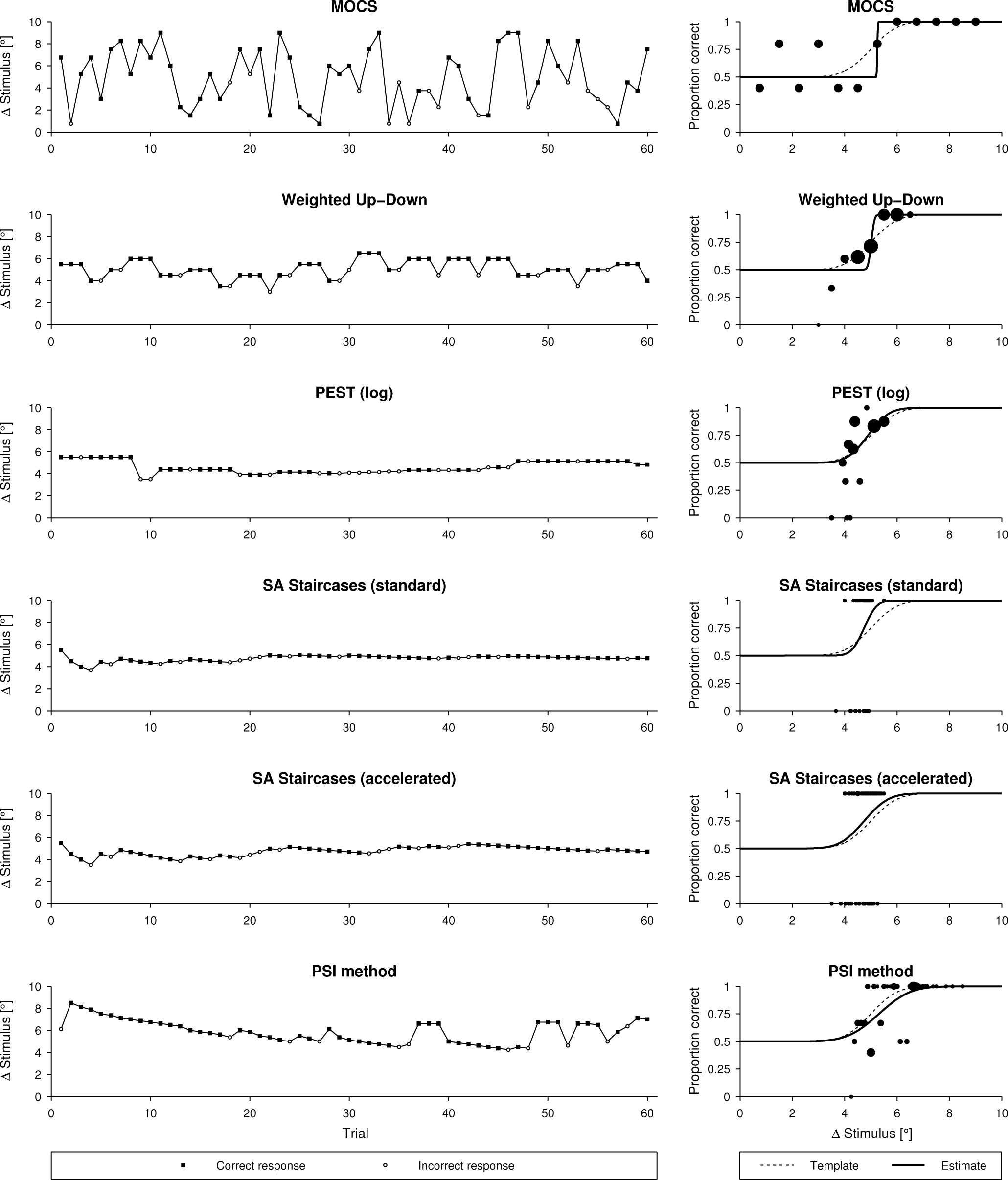
Examples of simulated sequences and psychometric functions for different sampling procedures. Examples for a template 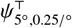 and comparison of the psychometric function of the template with the resulting fit using a Maximum Likelihood criterion. The size of the black dots indicates the number of repetitions at same stimulus level.

#### 4.3.2 Constant and variable errors of threshold and slope estimates

For each psychometric template and procedure, the *CE* and *VE* were calculated for the estimation of the threshold (Figure 6 and Figure 7) and the slope (Figure 8 and Figure 9). In these figures, it can be observed that for all sampling procedures except MOCS the absolute value of the threshold *CE* was below 0.1° for the largest part of the threshold and slope parameter space. For the PSI method, the *CE* was around 0.01°. In the case of MOCS and Weighted Up-Down, the threshold *CE* presented ripples depending on the threshold axis with absolute biases up to 0.4° and 0.2°, respectively. All methods showed decreasing performance towards the boundaries of the parameter space, especially for the smallest slopes. The standard SA Staircases showed a large negative bias for increasing thresholds and decreasing slopes (*CE* up to 1.5°) and large positive bias for low thresholds and small slopes (*CE* up to 1.2°). Similar but less pronounced effects could be found for the accelerated version of the SA Staircases. For the PSI method, the *CE* increased up to 0.8° for low thresholds.

**Figure 6.**
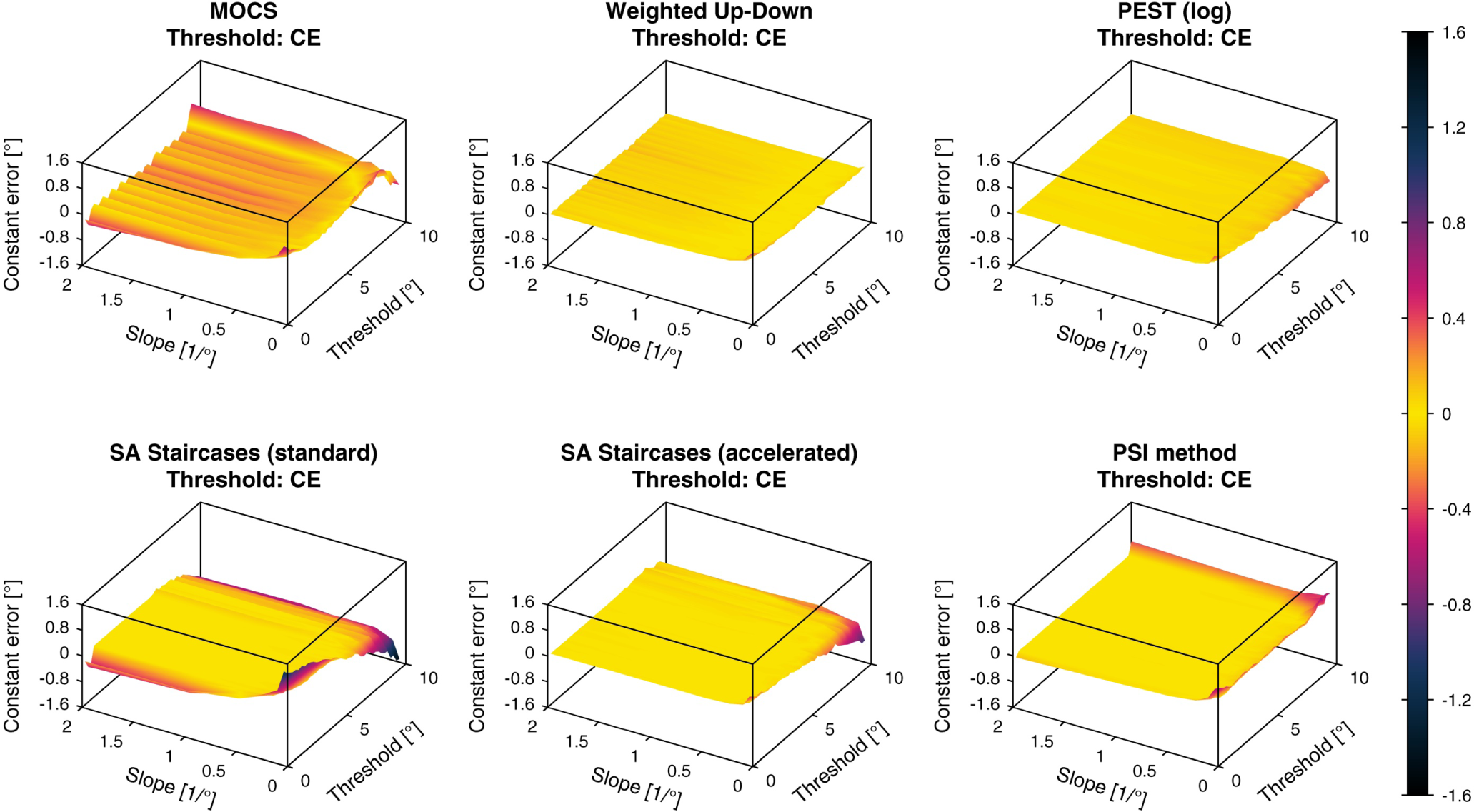
Constant errors of threshold estimates. Constant errors (*CE*) of the threshold estimates for the six different simulated sampling procedures. The color bar indicated the *CE* values in (°).

**Figure 7.**
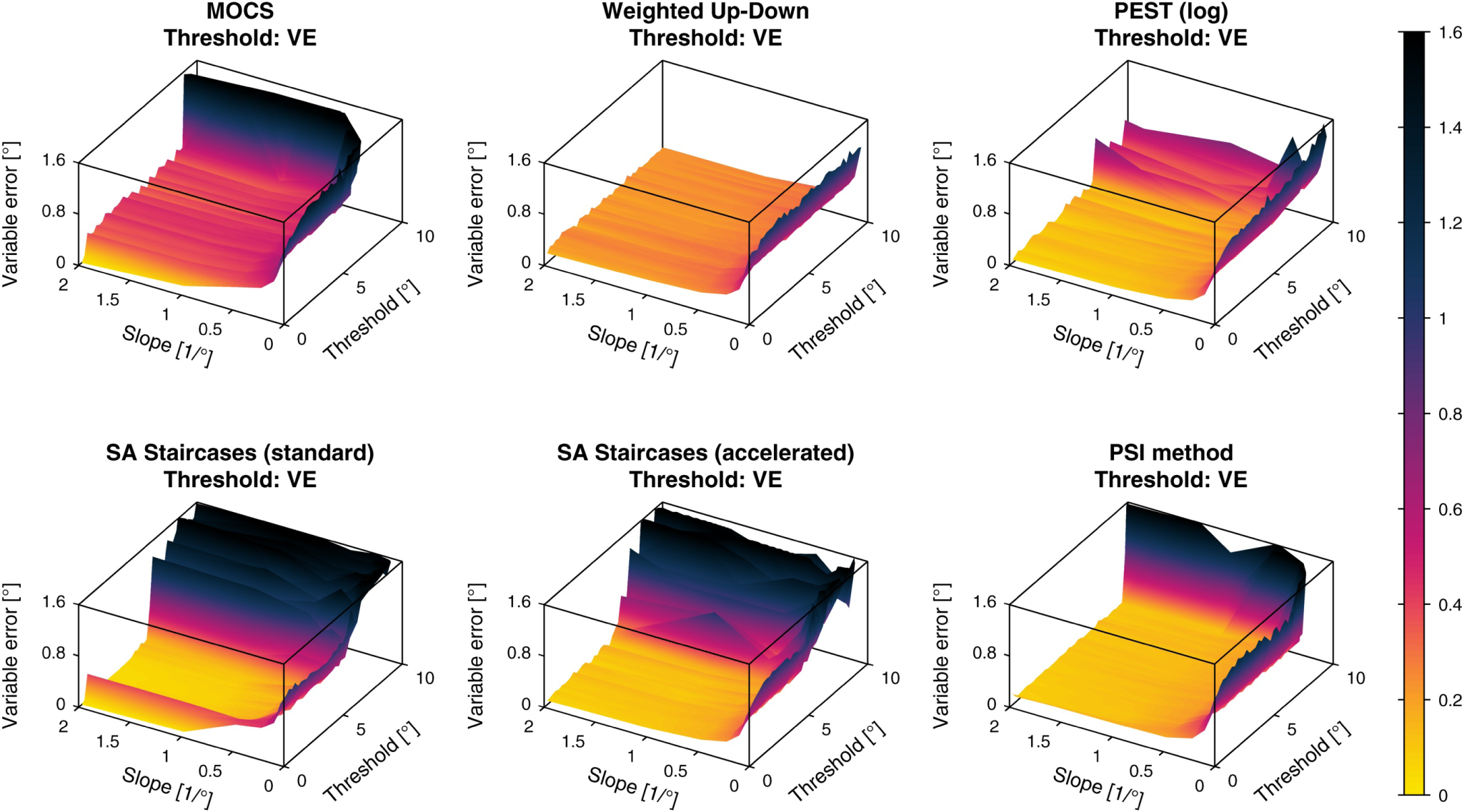
Variable errors of threshold estimates. Variable errors (*VE*) of the threshold estimates for the six different simulated sampling procedures. The color bar indicated the *VE* values in (°).

**Figure 8.**
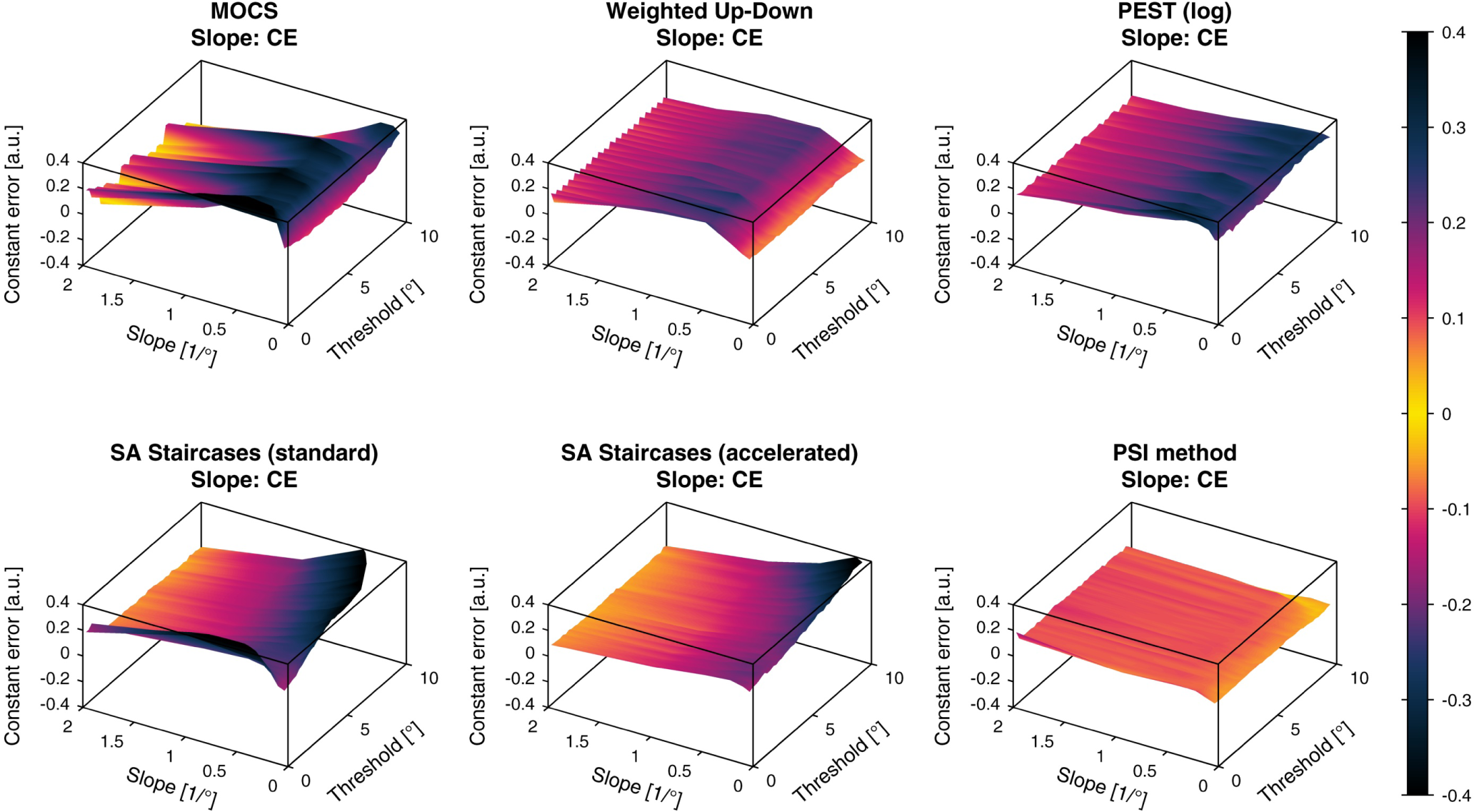
Constant errors of slope estimates. Constant errors (*CE*) of the slope estimates for the six different simulated sampling procedures. The color bar indicated the *CE* values in (a.u.).

**Figure 9.**
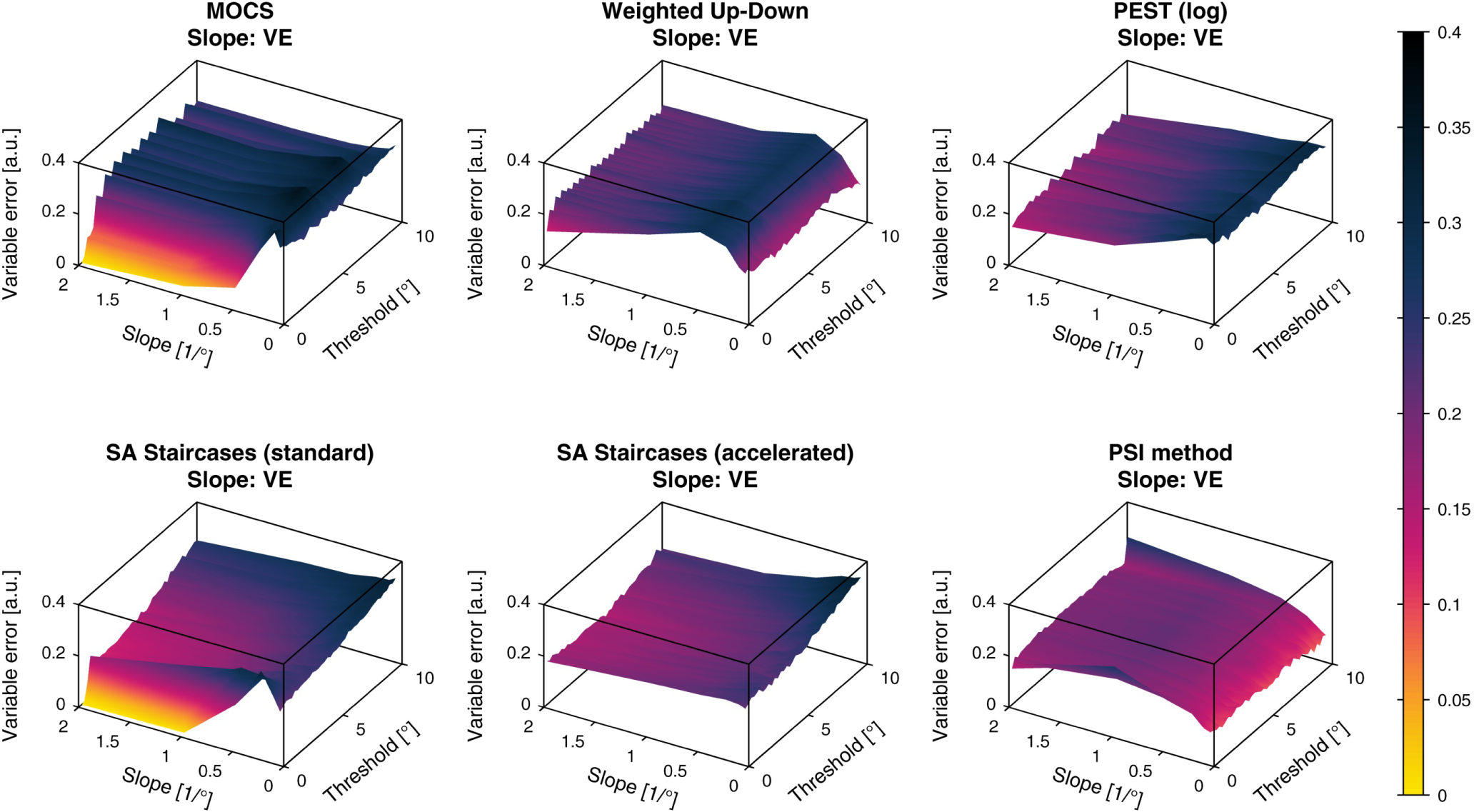
Variable errors of slope estimates. Variable errors (*VE*) of the slope estimates for the six different simulated sampling procedures. The color bar indicated the *VE* values in (a.u.).

The *VE* of the threshold showed much less consistent results compared to the *CE* with more oscillations along the threshold axis. The *VE* increased severely for small slopes in all methods. For thresholds larger than around 6°, *VE* started to increase non-monotonically to more than 1° for both standard and accelerated SA Staircases. For MOCS and the PSI method this was only the case for thresholds larger than 9°. The Weighted Up-Down method was the only sampling procedure not showing effects depending on the threshold value, neither for the central region of the parameter space, nor for the boundary. The PSI method showed the lowest *VE* of around 0.1° for almost the entire parameter space, compared to the other sampling procedures.

The slope *CE* showed an overestimation of the slope in the whole parameter space for all simulated sampling procedures. Similar to the threshold *CE*, slope *CE* ripples were found for MOCS and Weighted Up-Down. Both SA Staircases methods showed increasing bias for large thresholds and small slopes, whereas the standard SA Staircases also showed an increased bias for low thresholds. MOCS, Weighted Up-Down and PEST showed increasing overestimation for lower slopes, but bias decreased again for slopes smaller than 0.5/°. The most homogeneous *CE* across the parameter space (around 0.1) could be found for the PSI method and it was the only procedure for which bias decreased monotonously with smaller slopes. For illustration, a *CE* of 0.2 corresponds to an estimated slope of 1/° for a slope template of 0.5/°.

The slope *VE* showed similar behavior across the parameter space as the slope *CE*. For all methods the *VE* laid between 0.15 and 0.3. The main difference compared to the slope *CE* was the decreasing variability for small thresholds for MOCS and the standard SA Staircases.

#### 4.3.3 Percentage within bounds

The *PCTw*/*iB* curves are presented for each sampling procedure in Figure 10 for the threshold estimates and Figure 11 for the slope estimates. In general, the *PCTw*/*iB* curves for the threshold estimates followed the shape of exponential cumulative density functions. All methods had in common that for smaller slopes the *PCTw*/*iB* curve increases less rapidly, and the variability across thresholds was lower compared to the variability across different slopes. PEST, the PSI method, and the accelerated SA Staircases demonstrated the best performance and smallest variability across thresholds (e.g., reaching up to between 80% and 90% for a slope of 2/° and a threshold tolerance of ±0.1°). However, for slopes of 0.0625/° even these methods achieved only between 60% and 70% within a tolerance interval of ±1°. On the other hand, MOCS and standard SA Staircases showed the largest variability and lowest ratio of estimates for given tolerance bounds.

**Figure 10.**
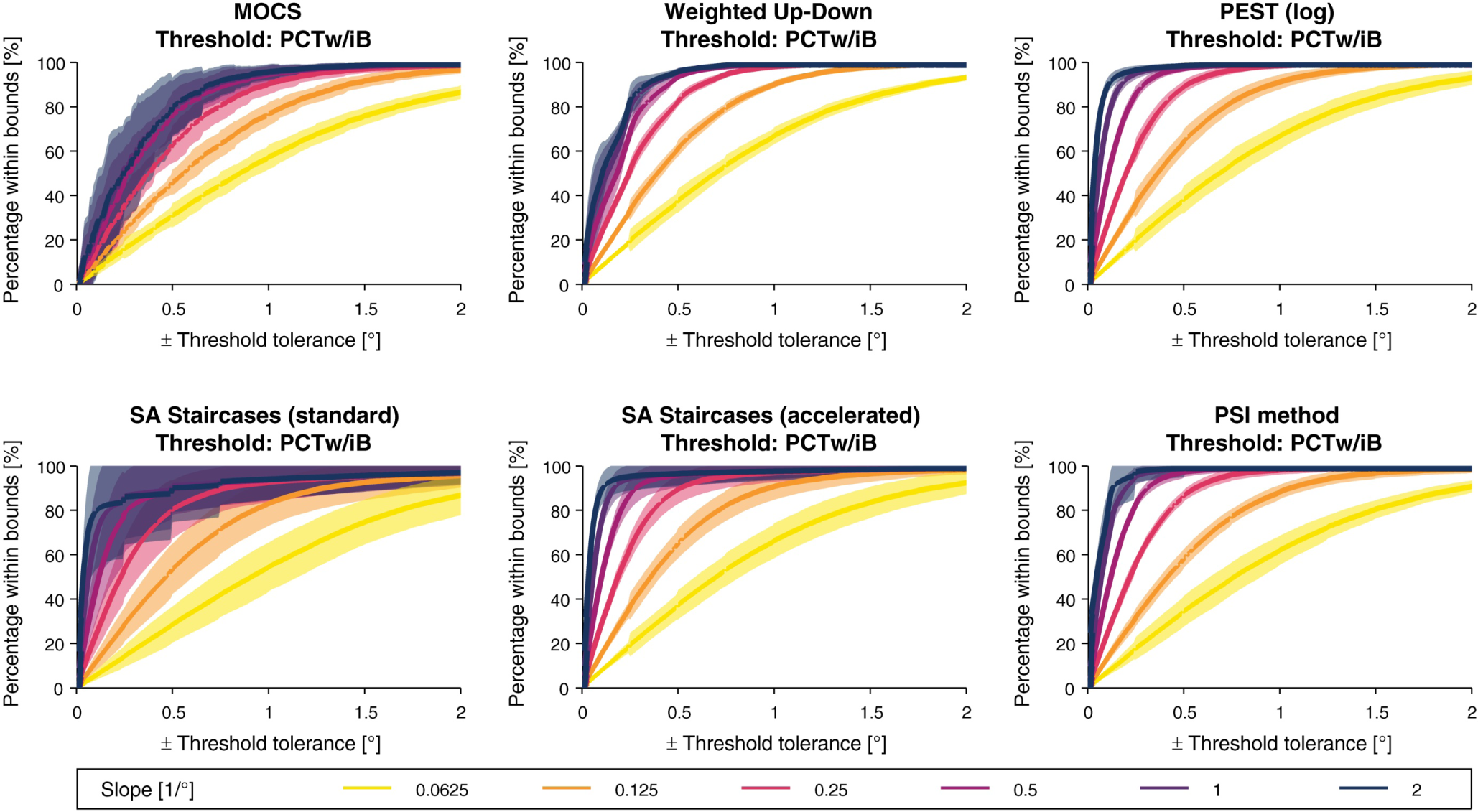
Percentage within bounds for the threshold estimates. For each slope value of the template parameter set, the *percentage within bounds* (*PCTw*/*iB*) function for threshold estimation are shown with the mean (bold line) and standard deviation (shaded band) across all modeled thresholds.

**Figure 11.**
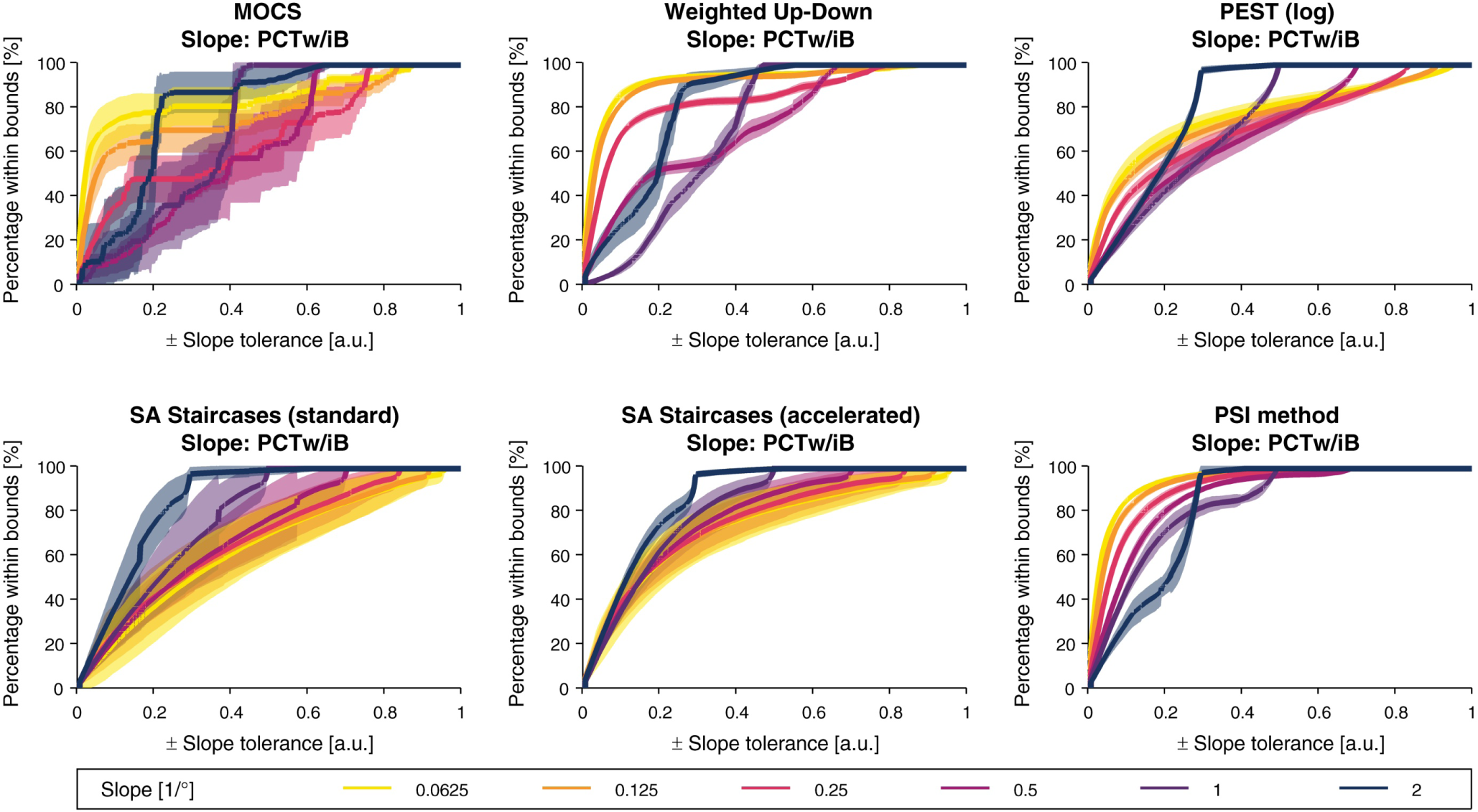
Percentage within bounds for the slope estimates. For each slope value of the template parameter set, the *percentage within bounds* (*PCTw*/*iB*) function for slope estimation are shown with the mean (bold line) and standard deviation (shaded band) across all modeled thresholds.

In contrast to the *PCTw*/*iB* curves for the threshold estimates, the derivative of the *PCTw*/*iB* curves did not monotonously decrease for the slope estimates. For all procedures except both SA Staircases, the *PCTw*/*iB* for small slope tolerance intervals was higher for smaller slopes compared to larger slopes, but was the other way around for larger intervals (i.e., larger than ±0.2). In the case of both SA Staircases, *PCTw*/*iB* was higher for larger slopes independently of the slope tolerance interval. The PSI method showed the best overall performance and reached 80% for all slopes for a tolerance of ±0.3. As for the threshold *PCTw*/*iB*, MOCS and standard SA Staircases showed the largest variability.

The tolerance-dependent overall *PCTw*/*iB* curves are presented for each sampling procedure in Figure 12 for the threshold estimates and Figure 13 for the slope estimates. For the threshold estimates, overall *PCTw*/*iB* reached 80% for all methods except MOCS for a tolerance interval of ±0.3°. PEST, accelerated SA Staircases, and the PSI method showed similar performance and outperform the others. The overall *PCTw*/*iB* for the slope was the highest for the PSI method and the accelerated SA Staircases, mostly independent of the slope tolerance interval size.

**Figure 12.**
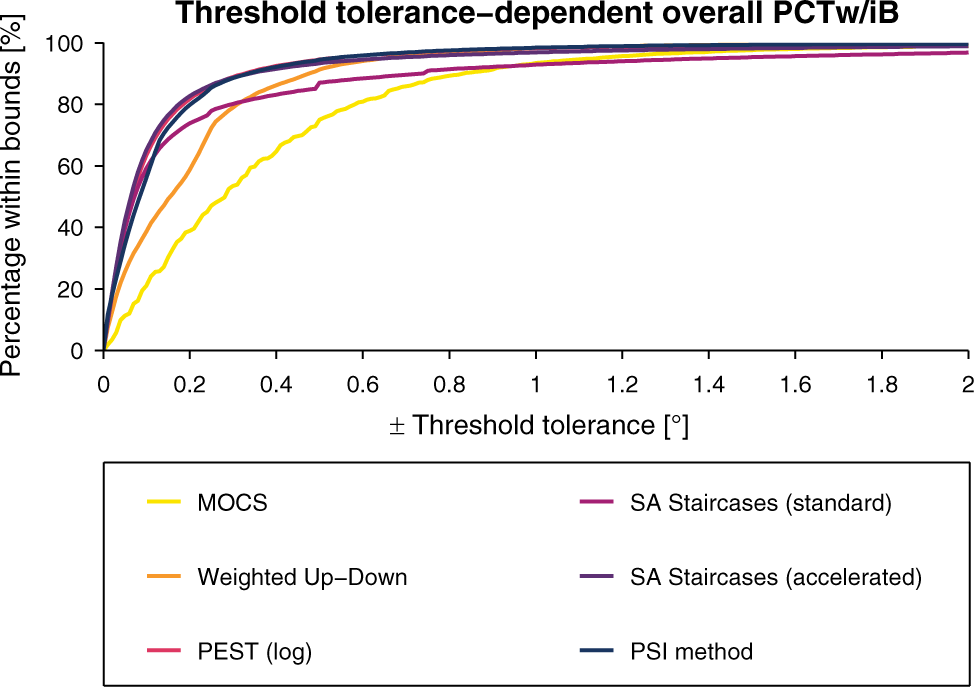
Overall percentage within bounds for the threshold estimates. Overall *percentage within bounds* (*PCTw*/*iB*) function for threshold estimation for a linearly resampled parameter space.

**Figure 13.**
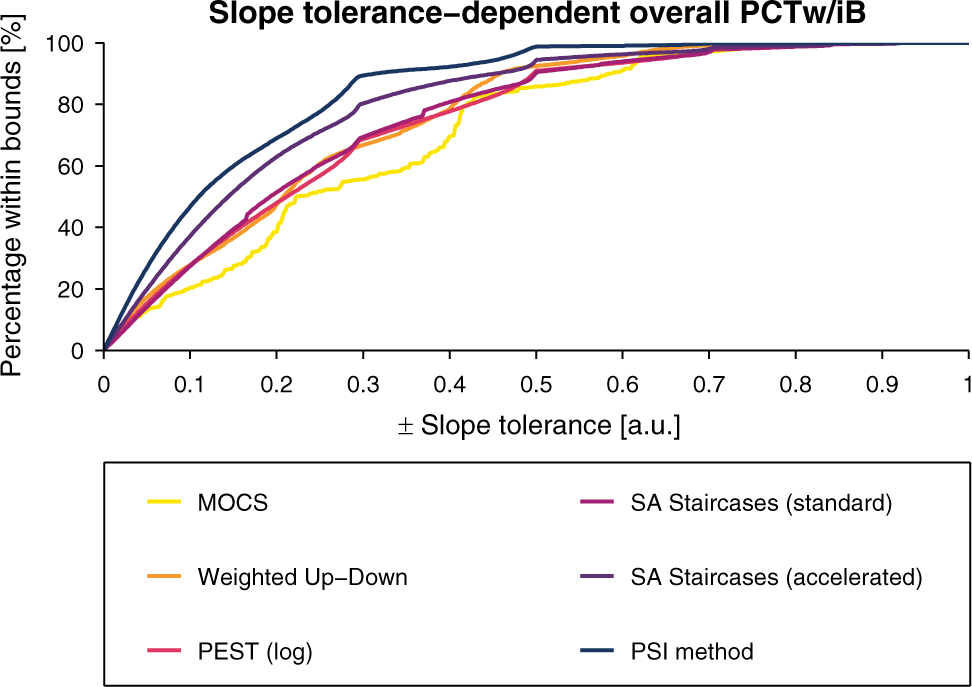
Overall percentage within bounds for the slope estimates. Overall *percentage within bounds* (*PCTw*/*iB*) function for slope estimation for a linearly resampled parameter space.

#### 4.3.4 Normalized area under the curve

The *nAUC* values computed individually for each combination of the two-dimensional parameter space are shown as surfaces for each sampling procedure in Figure 14 for the threshold estimates and Figure 15 for the slope estimates. These *nAUC* surfaces reflected the area under the *PCTw*/*iB* curves, showing a decrease of *nAUC* for smaller slopes for threshold estimates across all methods. In addition to Figure 10 the *nAUC* surfaces revealed threshold-dependent differences in performance, such as visible for both versions of the SA Staircases. For the slope estimates, both versions of the SA Staircases showed monotonously decreasing *nAUC* for smaller slopes, whereas in particular MOCS and Weighted Up-Down showed a large increase of performance for slopes below 0.5/°.

**Figure 14.**
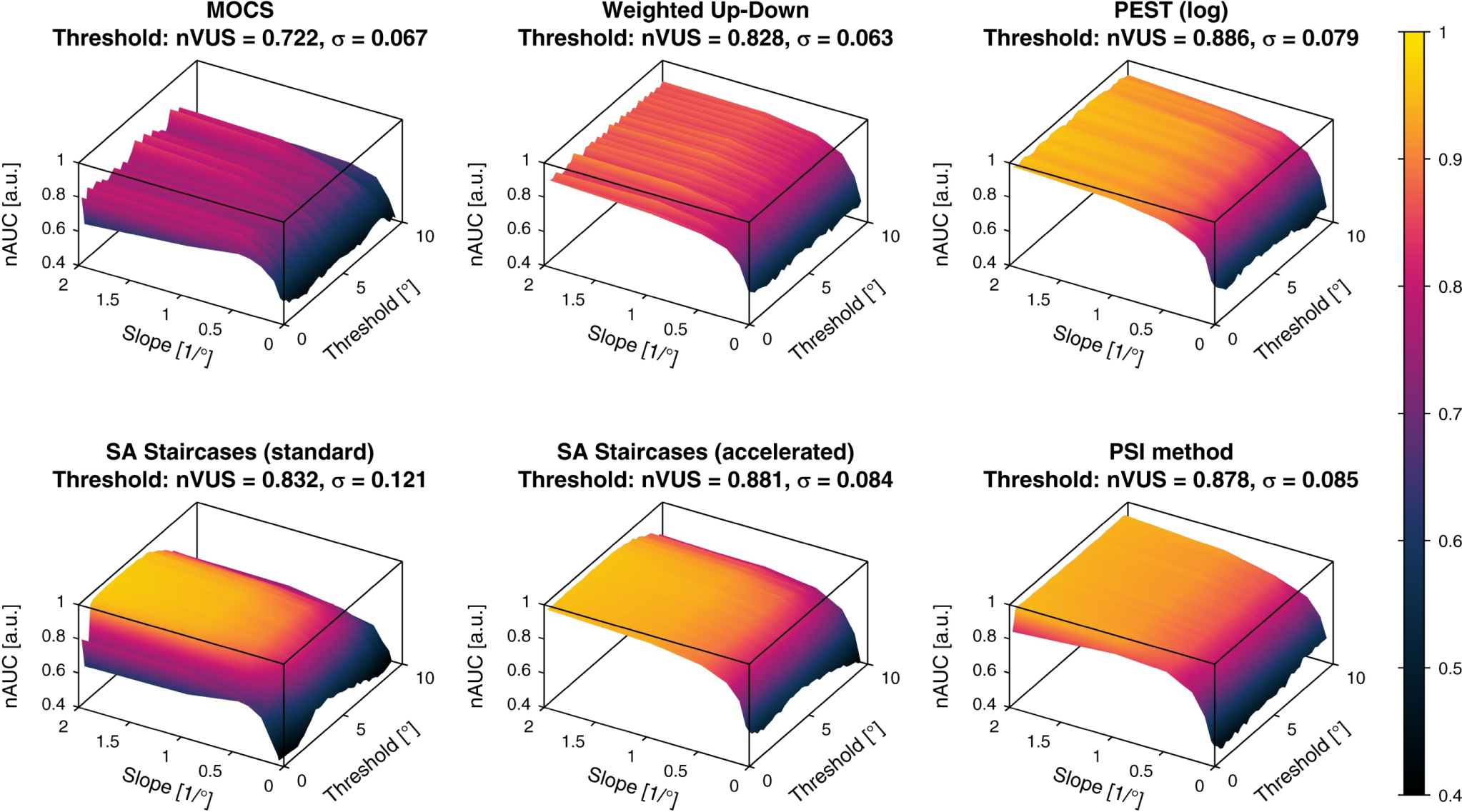
Normalized area under the curve for the threshold estimates. The *normalized area under the curve* (*nAUC*, represented by surfaces for the parameter space) and corresponding *normalized volume under the surface* (*nVUS*) and inhomogeneity parameter *σ* are shown for the threshold estimation. *nAUC* and *nVUS* values of 1 correspond to a perfect threshold estimation. The color bar indicates the *nAUC* values in (a.u.).

**Figure 15.**
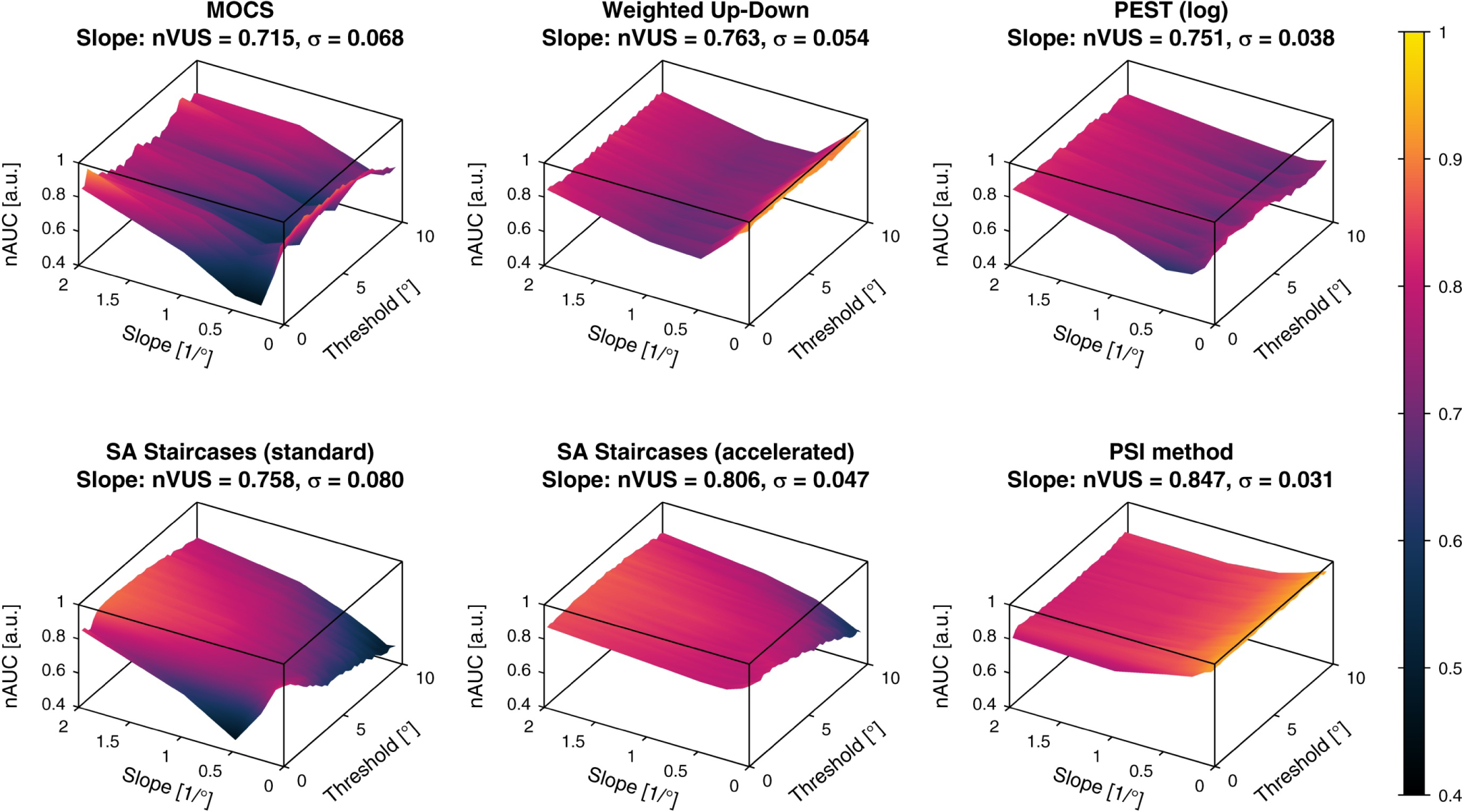
Normalized area under the curve for the slope estimates. The *normalized area under the curve* (*nAUC*, represented by surfaces for the parameter space) and corresponding *normalized volume under the surface* (*nVUS*) and inhomogeneity parameter *σ* are shown for the slope estimation. *nAUC* and *nVUS* values of 1 correspond to a perfect slope estimation. The color bar indicates the *nAUC* values in (a.u.).

#### 4.3.5 Normalized volume under the surface

The *nVUS* and inhomogeneity parameter *σ* are listed for each sampling procedure in Figure 14 for the threshold estimates and Figure 15 for the slope estimates. To compare the overall performance of the methods, these metrics are presented as ellipses (with center *nVUS* and half-axis *σ*, for both threshold and slope) in Figure 16. For the threshold estimates, PEST, accelerated SA Staircases, and the PSI method performed almost identically in *nVUS* (around 0.88) and *σ* (around 0.08). The overall best slope estimates were provided by the PSI method (*nVUS* = 0.85, *σ* = 0.03) followed by the accelerated SA Staircases. MOCS showed by far the worst performance for both threshold (*nVUS* = 0.72, *σ* = 0.07) and slope (*nVUS* = 0.72, *σ* = 0.07) estimates, and the standard SA Staircases showed the largest inhomogeneities (threshold: *σ* = 0.12, slope: *σ* = 0.08).

**Figure 16.**
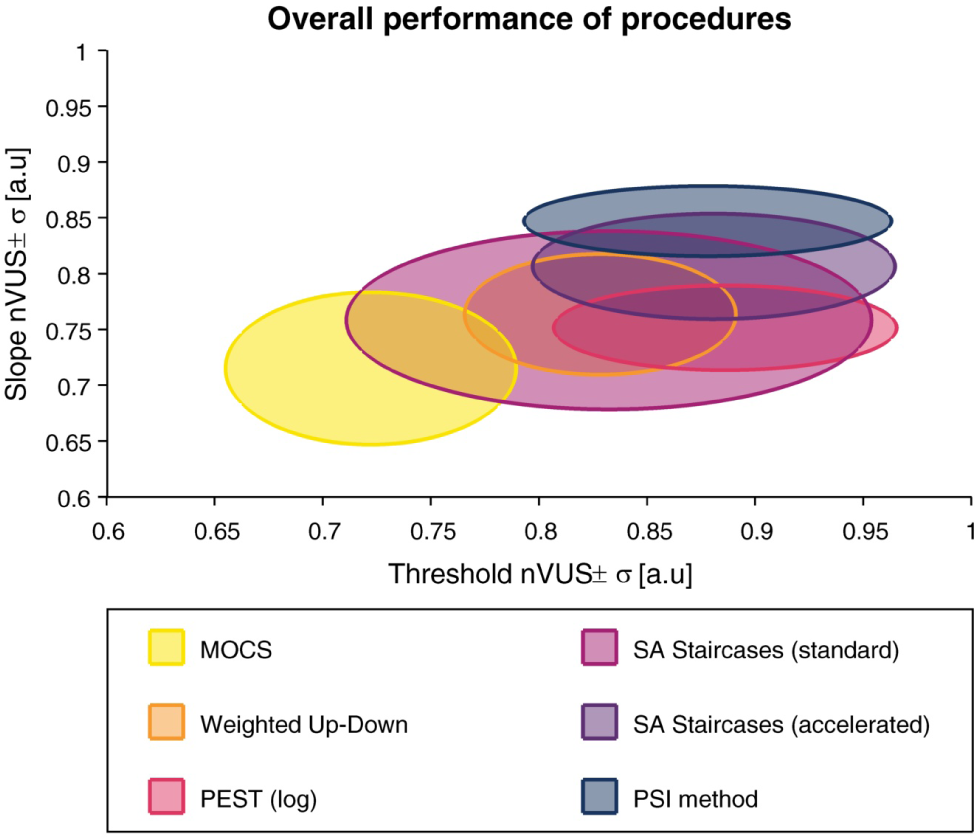
Overall performance of the six procedures. Two-dimensional plot showing the overall performance of the six methods based on the *normalized volume under the surface* (*nVUS*) and inhomogeneity parameter *σ* for the threshold and slope. The ellipses are centered at *nVUS* and the half-axes correspond to *σ*. Higher *nVUS* values and smaller half-axed indicate better estimation performance.

### 4.4 Discussion

The aim of this work was to introduce metrics to quantify the performance of psychophysical sampling procedures in application-driven simulations. The usefulness of the proposed metrics for in-depth analysis to identify how the procedures or their parameters could be tuned to potentially improve estimates and for choosing the best procedure for a specific application given some requirements (e.g., tolerance interval for the estimates) was illustrated using computer simulations. These included the simulation of six different sampling procedures were using values and ranges from a real-world application example.

#### 4.4.1 Strengths of this analysis framework

The efficiency in terms of variability multiplied by the number of trials (Taylor and Douglas Creelman, 1967; Taylor, 1971) has been evaluated for many sampling procedures using computer simulations (Taylor and Douglas Creelman, 1967; King-Smith et al., 1994; Watson and Fitzhugh, 1990; Hall, 1981; Findlay, 1978; Green, 1993; Madigan and Williams, 1987; Pentland, 1980; Simpson, 1989; Kaernbach, 1991). This approach aims at evaluating the benefit of adding more trials for better estimates. While this may be beneficial in some applications or for benchmarking, this analysis approach is incomplete and of limited use for real applications, where a sampling procedure should be selected and tuned to a specific distribution of psychometric functions of the population to be assessed. Furthermore, the performance of sampling procedures is often investigated for a single psychometric function or for a very limited number of threshold and slope parameters (Watson and Fitzhugh, 1990; Findlay, 1978; Green, 1993; Madigan and Williams, 1987; Pentland, 1980; Simpson, 1989; Faes et al., 2007; Kaernbach, 1991).

As shown in the present work, the performance of sampling procedures can vary considerably for different parameter values of psychometric functions. Therefore, results may not be representative for other values. As it has been noted by Klein: “*It is always a good idea to carry out Monte Carlo simulations of one’s experimental procedures, looking for the unexpected.*” (Klein, 2001). With the presented analysis method we could demonstrate the importance of simulating a wide range of psychometric functions covering the full parameter space of interest (i.e., threshold and slope) and illustrate the benefits of the different metrics for selecting and improving psychophysical sampling procedures.

The raw errors can describe bias (*CE*) and variability (*VE*). Their representation in the parameter space allows to detect suboptimal performance, for example, due to poor selection of parameters of a sampling procedure or decreases in performance towards the boundaries of the parameter space. By applying the transformation to arbitrary units on the slopes, errors in slope estimates can also be calculated and analyzed the same way, without penalizing slope errors in steep psychometric functions. For applications where a certain tolerance of estimation errors can be accepted, a new metric (*PCTw*/*iB*) based on the absolute errors (*AE*) was introduced. This metric is useful to assess the probability of the estimated parameter falling into a defined tolerance interval, and can be plotted for different psychometric functions. One advantage of the *PCTw*/*iB* compared to the average *CE* and *VE*, or the sweat factor (Taylor and Douglas Creelman, 1967; Taylor, 1971) is that this metric is robust against large outliers. As is the case in Figure 10, the plots can show very poor performance for small slopes. However, this visual representation of simulated templates may be misleading when deducing the overall performance. This has been addressed by post-hoc linear resampling of the parameter space of *PCTw*/*iB*_±*Tol*_ for each tolerance, from which the overall performance (tolerance-dependent overall *PCTw*/*iB*) can be calculated (Figure 12). This plot reveals that the overall performance for this parameter space is higher than what could have been misinterpreted based on Figure 10. Thus, despite large threshold estimate variability for small slopes, the best simulated methods provide around 80% of the threshold estimates within a tolerance interval ±0.2°, which in the present application is around one order of magnitude smaller than the proprioceptive difference thresholds of healthy subjects (ranging from around 1° to 5°) (Tan et al., 1994; Brewer et al., 2005; Tan et al., 2007; Lambercy et al., 2011; Rinderknecht et al., 2014; Elangovan et al., 2014; Cappello et al., 2015). The *PCTw*/*iB* is similar to the concept of a “usability index” introduced by García-Pérez and Alcalá-Quintana (2005), describing the percentage of times that a sampling procedure produces “usable data”. While authors present the “usability index” as a function of number of trials, our metric is a function of the acceptable tolerance and has a relation to the specific application examples. To deduce higher-level overall performance metrics, the *nAUC* surface (based on the integral of the normalized *PCTw*/*iB*) can be computed. Since the *nAUC* is remotely based on the *AE*, which is a composite of *CE* and *VE*, the *nAUC* surfaces are similar to the inverse of the *CE* and *VE* surfaces, but normalized to [0, 1]. Whereas the *nAUC* is calculated for each simulated psychometric function, the general performance metric (*nVUS*) and inhomogeneity parameter (*σ*) are independent of the parameter space, and can be used to select the optimal sampling procedure and to compare different sets of sampling procedure-specific parameters. The latter two metrics can, in case of threshold and slope to be estimated, be visualized as ellipses allowing a summarizing view on the performance for both parameters (Figure 16). Furthermore, this analysis framework is independent of the parametric version of the psychometric function and paradigm used, and can thus also be used, for example, for yes-no experiments with psychometric functions with γ = 0. Moreover, it can also be extended to the estimation of the lapse rate.

#### 4.4.2 Comparison of psychophysical sampling procedures for the application example

In this application example and given the chosen parameters of the sampling procedures, PEST provides the best overall threshold estimates (i.e., highest *nVUS*), as visible in Figure 16. Yet, the PSI method and the accelerated SA Staircases perform similarly well to PEST and distinctively better than the other sampling procedures. This can be confirmed in Figure 12 showing the tolerance-dependent estimation performance. Interestingly, the accelerated SA Staircases performs better than, for example, the Weighted Up-Down method, despite large *VE* and increased bias (*CE*) for high thresholds, whereas the latter shows more constant performance across the threshold parameter space (Figure 7). These high threshold *VE* oscillations (appearing for both SA Staircases) exist only for thresholds larger than the start level and may arise from the asymmetric descending and ascending step sizes. A similar, yet not so strong, effect can be noticed for PEST. Since with the logarithmic version of PEST steps become larger when ascending towards higher levels, it requires more trials to converge towards a threshold with the same precision. However, as the number of available trials is limited by administration time requirements of this application, variability of the estimates increases. Thus, a recommendation to improve the performance of these three procedures would be to select a higher start level and converge towards the threshold with descending steps. The reason for the Weighted Up-Down method not showing such decreases in performance for high thresholds, in contrast to the other adaptive methods, may be the constant step size which is well chosen with regards to the parameter space of the simulated templates. Unsurprisingly, the estimation performance of MOCS shows ripples in both *CE* and *VE* (Figure 6 and Figure 7). This reflects the limited number of different stimulus levels and presentations leading to equal estimations of psychometric functions for similar templates creating error ripples depending on the threshold. Both MOCS and the PSI method present increased *VE* at the boundary of the parameter space (i.e., high thresholds). This is attributable to the boundaries of the grid-like stimulus levels and the grid of psychometric functions used for the posterior probability distribution of the PSI method. As, for example, in the case of MOCS, the maximum stimulus level presented is 9°. If the threshold of the template to be estimated is higher than this maximum level, there is no data at levels with high performance available for the Maximum Likelihood fit of the psychometric function. Furthermore, each level is presented only five times, resulting in a discretization of the proportions of correct responses with steps of 0.2. This is not sensitive enough to properly estimate the psychometric function just based on data from low performance levels. Thus, to improve MOCS and the PSI method for this application, the threshold grids should be expanded beyond the parameter space, which would resolve the issues at the boundary. Across all sampling procedures, the threshold estimation performance decreases when the psychometric functions to be estimated have a small slope. This is an inherent problem of psychophysical assessments, as a slowly rising psychometric function increases uncertainty and thus introduces higher variability in answers. Moreover, it becomes more difficult for threshold tracking methods to converge towards the threshold. Therefore, all sampling procedures perform poorly for small slopes, as visible in Figure 10.

While PEST, accelerated SA Staircases, and the PSI method perform similarly well in estimating thresholds, the PSI method outperforms the other methods in estimating the slope of the psychometric function (Figure 16 and Figure 13). The PSI method was also in particular developed to estimate both threshold and slope (Kontsevich and Tyler, 1999). Thus, this sampling procedure places the stimulus levels in a way to optimize the estimation of both parameters (i.e., also at levels of high and low percentage of correct responses). This is also reflected in the “jumps” of the stimulus levels of the sequence example shown in Figure 5. In contrast, the other adaptive methods (in particular PEST and both SA Staircases) show faster asymptotic behavior towards the threshold to be estimated. The exploration of the stimulus levels with the latter sampling procedures depends solely on the selected start level and the initial step size, and is thus not optimized to cover a wide range of the psychometric function. The accelerated SA Staircases procedure is more aggressive than the standard version (because the denominator in the equation for calculating the new level is smaller, resulting in larger steps) and thus leads to more oscillations around the threshold, which helps to estimate the slope. Therefore, if the slope is of interest, a sampling procedure which places trials at well separated levels should be used (Klein, 2001). However, if the differences between the levels are too large in comparison to the slope (which can be the case for MOCS and the Weighted Up-Down method if sampling procedure-specific parameters are not well chosen), steep slopes are difficult to estimate because the sampling procedure may not place levels in the “slope region” of the psychometric function. In addition, due to its non-adaptiveness (i.e., predefined and fixed stimulus levels), MOCS places too many trials in regions of little information content (i.e., of very low and very high percentage of correct responses) (Figure 5). Besides the proportion of correct responses suffering from low resolution, these trials do not significantly contribute to the Maximum Likelihood fit of the psychometric function, as by definition of the psychometric function *ψ*(*x*) the asymptotic values are already defined (i.e., 0.5 and 1). Thus, depending on whether the slope of the psychometric function to be estimated is small or steep, non-adaptive and adaptive fixed-grid-level procedures (e.g., MOCS, Weighted Up-Down) may perform better than threshold tracking procedures (e.g., PEST, standard and accelerated SA Staircases), and vice versa, as can be inferred from Figure 11. The PSI method can also be considered an adaptive fixed-grid-level procedure. While the variability of the slope estimates improves for small slopes, the differences are small across the parameter space, as shown by the small value of the inhomogeneity metric *σ*. Similar to the threshold estimates, MOCS shows threshold-dependent ripples in the performance of the slope estimation, also resulting in large inhomogeneity, due to suboptimal and coarse distribution of stimuli.

Many of our findings are in line with the literature, though the results from the simulations are difficult to compare quantitatively due to a large number of different variants of sampling procedures, analysis methods mostly focusing on overall efficiency, and other sampling procedure-specific parameter values and applications. In the literature, most sampling procedures were compared to MOCS, and all discourage from using MOCS, as it is inefficient (i.e., suffers from higher variability) compared to adaptive methods (Watson and Fitzhugh, 1990; García-Pérez and Alcalá-Quintana, 2005; Turpin et al., 2010) and provides more biased results (Taylor et al., 1983). Especially in scenarios where there is no prior information about the parameters of the psychometric function to be estimated (which is often the case in clinical assessments with a large range of conditions), stimuli may (initially) be placed far from the region of interest. Methods using stochastic approximation have also been investigated, with the accelerated SA Staircase procedure showing better performance compared to the standard version and to fixed step size staircase methods (Faes et al., 2007). An additional advantage of the SA Staircase procedures is that no assumptions of shape or parameters of the psychometric function are required, and is thus less susceptible to parameter mismatches (Treutwein, 1995; Faes et al., 2007). The accelerated SA Staircase procedure was also shown to have a similar performance as mean-Bayesian methods, which seem to have near-optimal performance (Treutwein, 1995). As in the present work, it was suggested to use a clear suprathreshold initial stimulus intensity (i.e., level at high proportion of correct responses) and a large initial step size, which should lead to a good performance independent of the slope of the psychometric function (Faes et al., 2007). If the full shape of the psychometric function is desired, fast threshold tracking methods with decreasing step sizes should be avoided (García-Pérez and Alcalá-Quintana, 2005). This could also be shown in the present simulations, although it is mostly valid for small slopes only.

#### 4.4.3 Limitations

One limitation of this analysis framework is the need to representatively sample the parameter space, in order to acquire enough information to see the behavior of different sampling procedures (e.g., threshold-dependent ripples induced by MOCS). This requires large computational processing power. However with today’s computers and multiprocessor architectures, the simulations can be parallelized for faster run time (e.g., to a few hours to a couple of days).

Since the shape of the psychometric function may not follow the mathematical model *ψ*(*x*) used when fitting the psychometric function, this might introduce systematic errors. However, this is a general concern of hybrid procedures (using heuristic adaptive sampling procedures combined with fitting of parametric psychometric functions) and methods assuming a particular shape for the stimulus selection. Nevertheless, the presented metrics could also be used with estimates resulting directly from the adaptive sequence instead of estimates from the fitting process. Depending on the perception modality assessed and the sampling procedure, these two types of estimates may highly correlate (e.g., (Rinderknecht et al., 2014)).

Since the performance metrics are intended to be used for an application-driven evaluation of sampling procedures, the present quantitative illustration is limited to the example of proprioceptive function, and a non-exhaustive set of different sampling procedures and method parameters was presented. Furthermore, to simplify the simulations and the presentation of the results, the guess rate γ = 0.5 and lapse rate *λ* = 0 were held constant for the templates as well as for the fitting process instead of using free parameters. Since it has been shown that not taking lapses into account may bias the threshold estimates (Wichmann and Hill, 2001a), and that perception can be non-stationary during experiments due to, for example, inattention, learning, or change in decision criteria (Watson, 1980; Hall, 1983; Leek et al., 1991; Cohen and Maunsell, 2011; Doll et al., 2015), more realistic simulations should take these factors into account, and more elaborate sampling procedures (Prins, 2013) or methods to address these specific challenges could be used in combination with the fitting process or the sampling procedure (Wichmann and Hill, 2001a; Rinderknecht et al., 2018). Since the perception thresholds and slopes of the sampled population may not be linearly distributed, as assumed in this work when resampling the *PCTw*/*iB*_±*Tol*_ or *nAUC* space, a more realistic distribution (e.g., log–normal distribution for parameters with semi-infinite positive support) could be based on some prior knowledge. This could be achieved by defining an n-dimensional (in the present case two-dimensional for the threshold and slope parameters) density function in the parameter space to correct *nVUS* with a weighting factor (e.g. multiplying the volume sections by a factor) or to calculate the inhomogeneity parameter *σ* based on a non-linear resampled space (i.e., resampling density according to the population distribution).

## 5 Conclusions

This work introduces a set of novel metrics to evaluate the performance of psychophysical sampling procedures in estimating parameters of psychometric functions quantitatively, using application-driven simulations. The analysis framework can be used for any type of sampling procedure and parameters to be estimated (e.g., threshold, slope, lapse rate), and is independent of the parametric versions of psychometric functions (e.g., normal, logistic, Weibull) and the application-specific parameter spaces. In summary, the illustrative analysis of a simulation based on a scenario of proprioceptive assessment using these metrics allowed identifying suboptimal parameter choices for different simulated sampling procedures and deriving suggestions on how to improve the methods. Furthermore, the optimal sampling procedure could be identified, which could be tuned and analyzed in a second iterative step using the same metrics framework. Thus, these metrics allow a deeper understanding of the strengths and limitations of the sampling procedures, facilitate the parameter tuning, and showed that it is important to evaluate the procedures for different psychometric functions with metrics beyond efficiency.

## Author contributions

MR, OL, and RG contributed to the conception of this work. MR developed the methodology, implemented the computer simulations, performed the analysis, interpreted the results, and drafted the manuscript. MR, OL, and RG revised the manuscript and approved the final version.

## Acknowledgments

The authors would like to thank K. Leuenberger and W. L. Popp for fruitful discussions. This research was supported by the ETH Zurich Foundation in collaboration with Hocoma AG, the Janggen-Pöhn Foundation, and the Swiss National Science Foundation through project 320030L_170163.

